# Green Index: a widely accessible method to quantify greenness of photosynthetic organisms

**DOI:** 10.1101/2023.08.23.554481

**Authors:** Santiago Signorelli, Esteban Casaretto, A. Harvey Millar

## Abstract

Plant phenotyping involves the quantitative determination of complex plant traits using image analysis. One important parameter is how green plant tissues appear to the observer, which is indicative of their health and developmental stage. Various formulas have been developed to quantify this by calculating leaf greenness scores. We have developed a revised formula called “Green Index” (GI) devised out of the need to quantitatively assess how the apparent greenness of seedlings changes during de-etiolation. The GI calculation is simple, uses widely available RGB values of pixels in images as input, and does not require commercial software platforms or advanced computational skills. In this study we describe the conception of the GI formula, compare it with other widely used greenness formulas, and test its wider application in plant phenotyping using the open source free software platform RawTherapee. We demonstrate the utility of the GI in addressing common issues encountered in assessing plant biology experiments, underscoring its potential as a reliable and accessible tool. Finally, we explore the correlation between GI and chlorophyll content, assess its reliance on different types of photography, and summarizes the key steps for its effective utilization.

## Introduction

Plant phenotyping focuses on the quantitative determination of complex plant traits such as anatomical, physiological and biochemical properties (Fiorani and Schurr 2013). The greater accessibility for image capture and image analysis software have contributed to the large expansion of non-destructive plant phenotyping pipelines in the last decade (Furbank and Tester 2011; Tao et al. 2022; Zavafer et al. 2023). Leaf colour is an important feature to capture in plant phenotyping. The colour of a leaf is often indicative of plant nutrition and the absence of abiotic and biotic stress and the greener a plant appears typically indicates the healthier a plant in perceived to be by the observer (Liang et al. 2017).

Human perceptions of colours are complex (Webster 2009) and a range of different variables have been developed to quantify them. The Munsell colours system was defined in 1911 (Burchett 2002) and adopted by the USDA as the official system to define colours in agriculture (Nickerson 1946). This system defines colours in three dimensions: value, hue, and chroma. Munsell hue (or colour hue) sets the colour as red, green, blue, or something in between (Cooper 1929). Munsell value indicates the lightness or darkness of the colour, ranged from 0 (pure black) to 10 (pure white) (Cooper 1929). Munsell chroma refers to the colour saturation, which can be weak (low chroma) or saturated (high chroma) (Cooper 1929). In plants, the hue colour of leaves is mainly determined by the absorption of chlorophylls, which determine a specific hue colour. However, under certain circumstances, the hue colour can be shifted to the green-yellow, if for example, the concentrations of chlorophylls are reduced but carotenoids are kept constant. When both chlorophyll and carotenoids are reduced in a same ratio, the colour hue is expected to be constant but the chroma will be reduced. In contrast, if both chlorophyll content and its ratio to other pigments are kept constant, but the leaf is observed under different light intensity, then the Munsell value varies, being 10 in complete darkness. Despite the utility of the Munsell system, it is not the colour system that dominates in digital images, instead digital images quantify colours using RGB, CMYK, HSL, HSV and PMS standards for a variety of technological reasons.

Like Munsell, some of these systems define colours based on variables such as hue, saturation, lightness, value or brightness. A notable except is RGB that does not define brightness (Munsell’s value), and thus the same RGB values can result in different colours on different devices. Instead, RGB define colours by the simple addition of the three primary colours (Red Green and Blue). RGB generally represents what is seen by the human-eye, as the three types of humans-cone are sensitive mainly to reddish, greenish, and bluish thirds of the visible spectrum (Trussell et al. 2005; Hunt and Pointer 2011). But it is widely acknowledged that the RGB color gamut is restricted in the blue-green colours, often leading to professional printers preferring CMYK (McGavin et al. 2005). Despite these limitations in blue-green representation, today it is RGB that dominates as a standard in digital images, for the Internet, most computers, printers, and many low-to medium-end consumer digital cameras and scanners (McGavin et al. 2005). This followed RGB cooperative creation and adoption by HP and Microsoft in 1996 (Stokes et al. 1996) and its standardization by the International Electrotechnical Commission (IEC) in 1999 (International Electrotechnical Commission 1999).

A series of different plant phenotyping studies have used colour analysis algorithms to calculate the proportion of green pixels in a given image to either determine plant canopy area or height e.g. (Starý et al. 2020; Abutaleb et al. 2021; Valluvan et al. 2023), or to calculate the proportion of green pixels in a given image of plants e.g. (Liang et al. 2017; Li et al. 2021). In some of these reports green pixel proportion correlated with chlorophyll content or photosynthetic rate. While other studies have aimed to score the greenness of a specific area based on the RGB values of each pixel (Gobron et al. 2000). This information can be acquired from conventional photos of plants grown in lab conditions or aerial photos of the canopy of plants in fields. This has the potential to evaluate changes in greenness associated with pathogen infection, environmental stresses, nutrient deficiency, and senescence. There are multiple formulas available to calculate greenness scores based on RGB values such as:

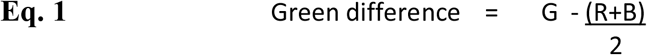

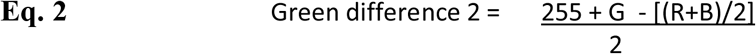

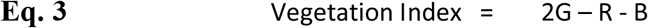

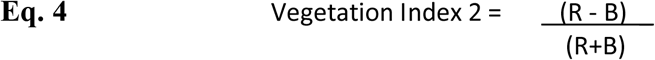

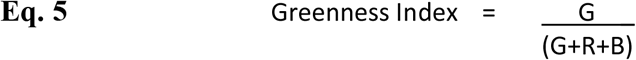

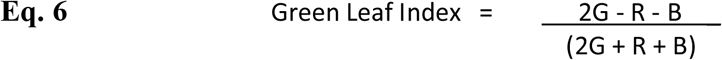

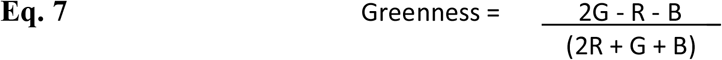

The Green Difference formulas (Eq. 1 and 2) are often used in photography to identify green pixels. Green Difference 2 (Eq. 2) is a variant of Green Difference (Eq. 1) that is aimed to avoid negative values. The Vegetation Index (Eq. 3) was developed to focus on green, identify the leaf area on a picture and contrast it against a non-green background (e.g. soil) (Woebbecke et al. 1995). Vegetation Index 2 (Eq. 4, Kawashima and Nakatani (1998)) was developed to estimate chlorophyll content, and compared to other formulas tested in their study, it was the index showing the best negative correlation with chlorophyll content. For simplicity of comparison we invert the conditional formatting for this formula to correlate better with other greenness indices. Greenness Index (Eq. 5), also known as Normalized Greenness Intensity (NGI), is another formula developed with the same propose (Yuan et al. 2017). Other greenness indexes referred to in the literature measure the green area over a total area in view, which is different to the Greenness Index we refer to here (Eq. 5). Green Leaf Index (Eq. 6) (Gobron et al. 2000; Javornik et al. 2023) is a widely used formula with the same aim, which has been incorporated into commercial phenotyping platforms such as the multispectral 3D scanner PlantEye F500 (Phenospex, Heerlen, Netherlands). Greenness (Eq. 7, Lazarević et al. (2022)), is slightly different to Green Leaf Index and is used by commercial phenotyping systems (e.g. Phenospex) under the name Green Leaf Index.

Recently, we developed a different formula based on RGB values we called “Green Index” (GI), to quantitatively discriminate between the degrees of greenness of arabidopsis seedlings undergoing de-etiolation in the light (Wijerathna-Yapa et al. 2021). We did this because the equations outlined above to obtain a greenness score could not discriminate the differences that appeared evident to us by eye. The GI calculations we developed did not require specific software or advanced skills. In this explanation of the method, we explain the development of GI formula, compare it to other popular methods and describe its usage with the open software platform RawTherapee in a way that is highly accessible without cost or computational skills. We also expanded its usage to consider application to other common problems presented in plant biology to validate the index and evidence its potential. We also test its correlation with chlorophyll content, evaluate its dependence on different types of photographic camera, and summarise the key steps for practical usage of GI.

### Experimental Procedures

#### Image acquisition

Pictures of 24-coloured card and plants were taken using a Nikon D7000 digital camera on a photographic stand supplemented with artificial light that were constant for all images produced by us. These pictures were used to develop the GI formula. For comparison of GIes obtained with a different camera, we also used an iPhone Xs smartphone. To test the GI in leaves subjected to different treatments, we used leaves or whole plant images from different publications and cited them accordingly. We used Affinity Photo and Affinity Designer V1 and V2 for figure preparation with colour format RGB/8 and colour profile wide Gamut RGB.

### Colour threshold for GI formula development

For developing the GI formula, we first analysed the picture or a 24-colour card using the Colour Thresholder app, within the Image Processing Toolbox MathWorks, applied it to RGB channels to define the range of RGB values excluding all colours but greens that can be found in leaves (a pale and a darker green) (see Supplemental Video 1 and Supp. Fig1D), and secondly, we defined the specific values within those ranges that gave the highest score for green. Likewise, we applied the same RGB thresholds to photos of arabidopsis (*Arabidopsis thaliana*) seedlings on agar plates confirming that those RGB thresholds exclude all the pixels of the picture except the leaves themselves. With this input, we proceeded to the mathematical development of the formula as detailed in the Results section.

### RGB determination

During formula development, RGB values were obtained using Affinity Photo and the Colour Picker tool. For the rest of the study, we used the RawTherapee programme as described in the Results section. RGB values vary between different programs (or even within the same program) if they are opened using different colour format and profile. Therefore, we ensured consistent colour format was used when opening multiple picture files.

## Results

### Green Index formula development

RGB values refers to the levels of red (R), green (G) and blue (B) in a coloured pixel and have numerical ranges between 0 and 255. Therefore, any coloured photo can be expressed as RGB and the RGB values for each pixel of the photo can be obtained with many basic image processing programmes because they are the IEC standard for colour in digital images (International Electrotechnical Commission 1999). However, anecdotally we found that Eq 1-7 did not necessarily discriminated what we observed as different levels of greenness by eye. As an initial approach to improve a formula, a term we called pre-GI_a was calculated as a number for which the R and B components of a pixel decreased the value whereas the G component value gave a maximum when it was 100, and the values were divided by 255 for each component to get a pre-GI_a value that sits always between 0 and 1, as follows:

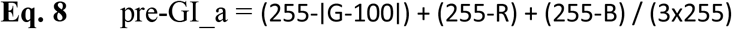

A picture of a 24-colour card was used to collect RGB values of each colour hue (**Figure S1A**) to test this preliminary formula. We observed that the pre-GI_a of a black square was high and the difference between green and some other colour hues was poor (**Figure S1B**). To better define which RGB values should score higher, we analysed the 24-colour card with the Colour Thresholder app from MATLAB®. By adjusting the different windows of each colour channel, we noticed that values from 75 to 255 for G and 0 to 75 for R and B allowed us to selectively pick greens hues excluding other colours (**Figure S1C-D**). The **Supplemental Video** explains this process. Based on this observation, we concluded that values around 165 (middle point between 75 to 255) should score the most for G and around 37.5 (middle point between 0 and 75) should score the most for R and B. We introduced this consideration in an improved version of the formula as follows:

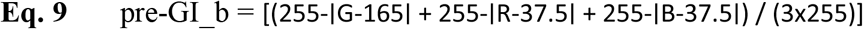

The new formula, pre-GI_b gave the highest scores to the different green colour hues tested (Dark Green and Green both getting values close to 1 (**Figure S1B**). To maximise the difference between green hues and any other colour, we divided Eq. 9 by the difference between 1 and the results of Eq. 9, as follows to generate pre-GI_c:

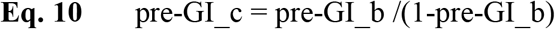

Eq. 10 implies that the index of Eq. 9 is divided by a small number for green and greater numbers for a colour different to green, thus maximising the differences between colours as observed in **Supp. Figure 1B**. Due to this addition to the formula, the results are no longer within the range of 0 and 1, and can yield very high values (e.g. when divided by a number near 0). We observed values as high as 11.8 from the 24-colour card. We therefore decided to normalize the data by dividing Eq. 10 by 12 to get all the values between 0 and 1, to generate a final GI as follows:

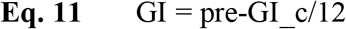

The results of analysing a 24-colour card suggest that the formula in Eq. 11 discriminates very well between different green values and saturations and any other colour hue.

Eq. 12 represents this GI formula independent of its derivation:

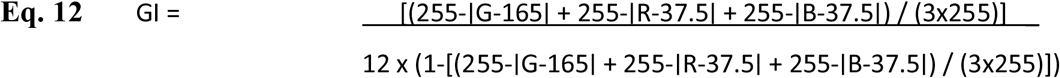

In a spreadsheet program such as Excel® the formula can be written as:

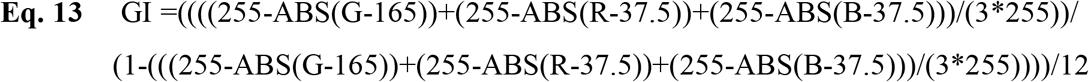

where the ABS function is used to indicate absolute value, and G, R, and B should be replaced by the spreadsheet cells containing 0-255 values for each colour. We then systematically compared the results of the GI formula with the results obtained with other formulas based on RGB values (Eq. 1 to Eq. 7). None of the other methods provide good correlations with green value and saturation and, in many cases, colour hues different to green are given a higher score than green (**Figure 1**).

**Figure 1.**
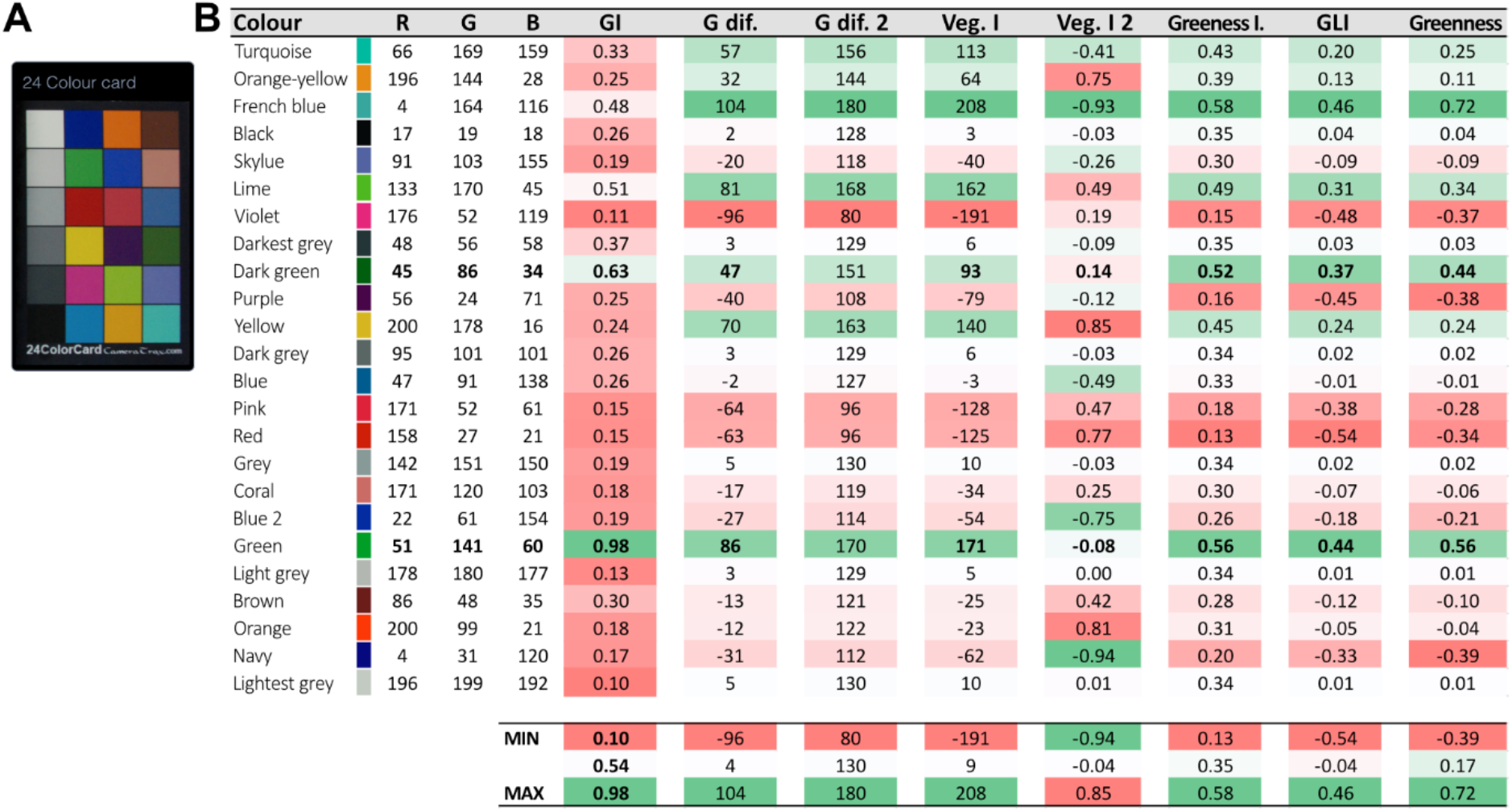
Analysis of the scores obtained by multiple methods based on RGB values in a 24-colour card. **A**. 24 colour card used in this study. **B**. RGB values for each colour and the scores obtained with the different formulas presented in introduction. Red fills indicate bad scores, whereas green fills indicate good scores.

### Extracting RGB values from plant images

A range of programs including Photoshop or Affinity Photo allow users to obtain RGB values with one click; however, for this study we chose to describe the process using the open source software RawTherapee (rawtherapee.com) which is available across all computer operating systems. In **Figure 2**, we show the extraction of RGB values for an example analysed in this study (**Figure 3**). We opened the image of interest with RawTherapee, selected the Lockable Colour Picker tool and clicked in 10 different areas of each leaf/treatment to get n=10. All the R, G, and B values were recorded in a table for later processing using the formula described above for GI (Eq. 13).

**Figure 2.**
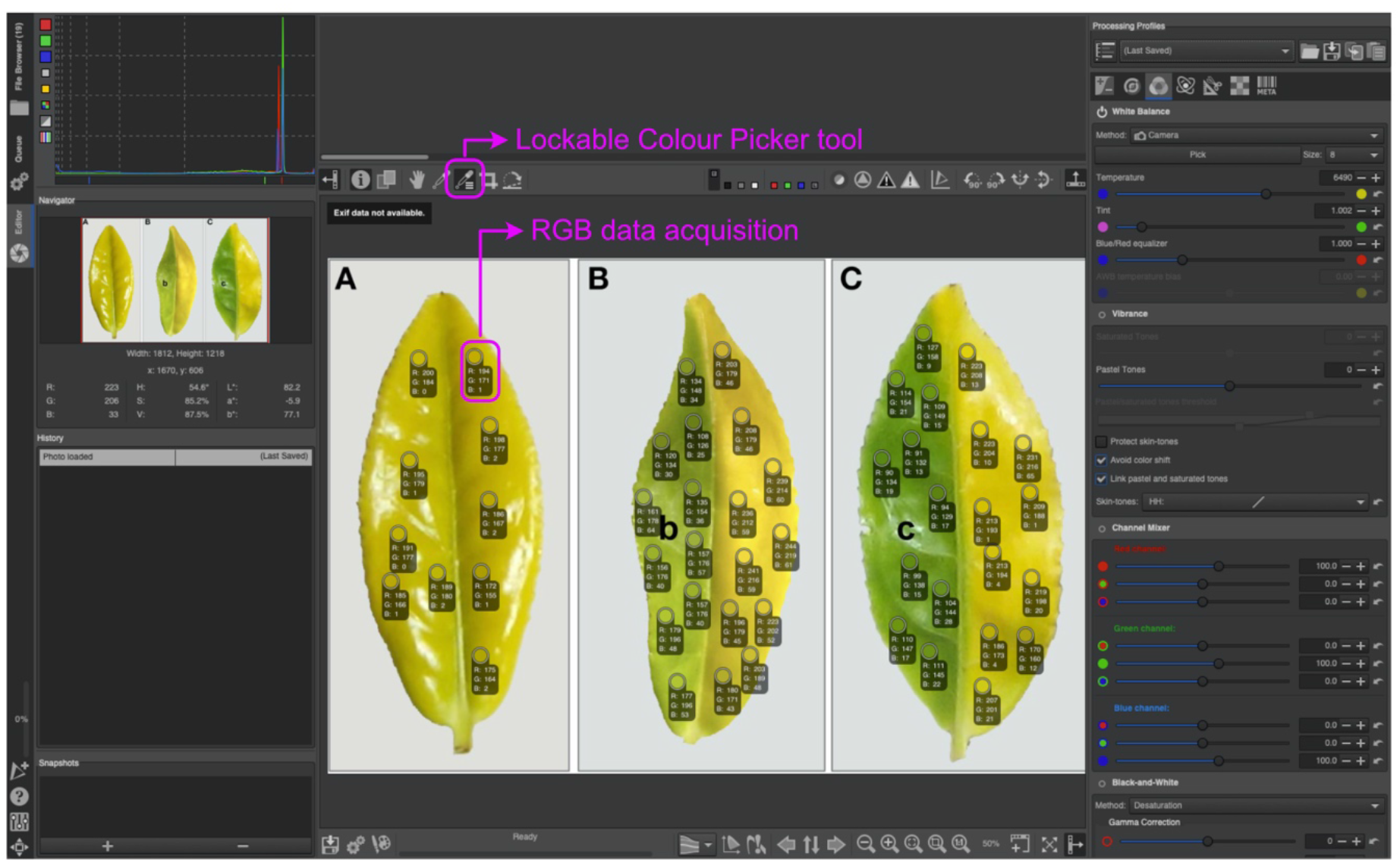
Collecting RGB values form an image using the open access software RawTherapee. The figure shows the Lockable Colour Picker tool, which can be used to select an area of interest in an image to obtain the RGB values. Multiple examples of the RGB values obtained for different areas of interest are recommended. Here 10 technical replicates were used per leaf/treatment.

**Figure 3.**
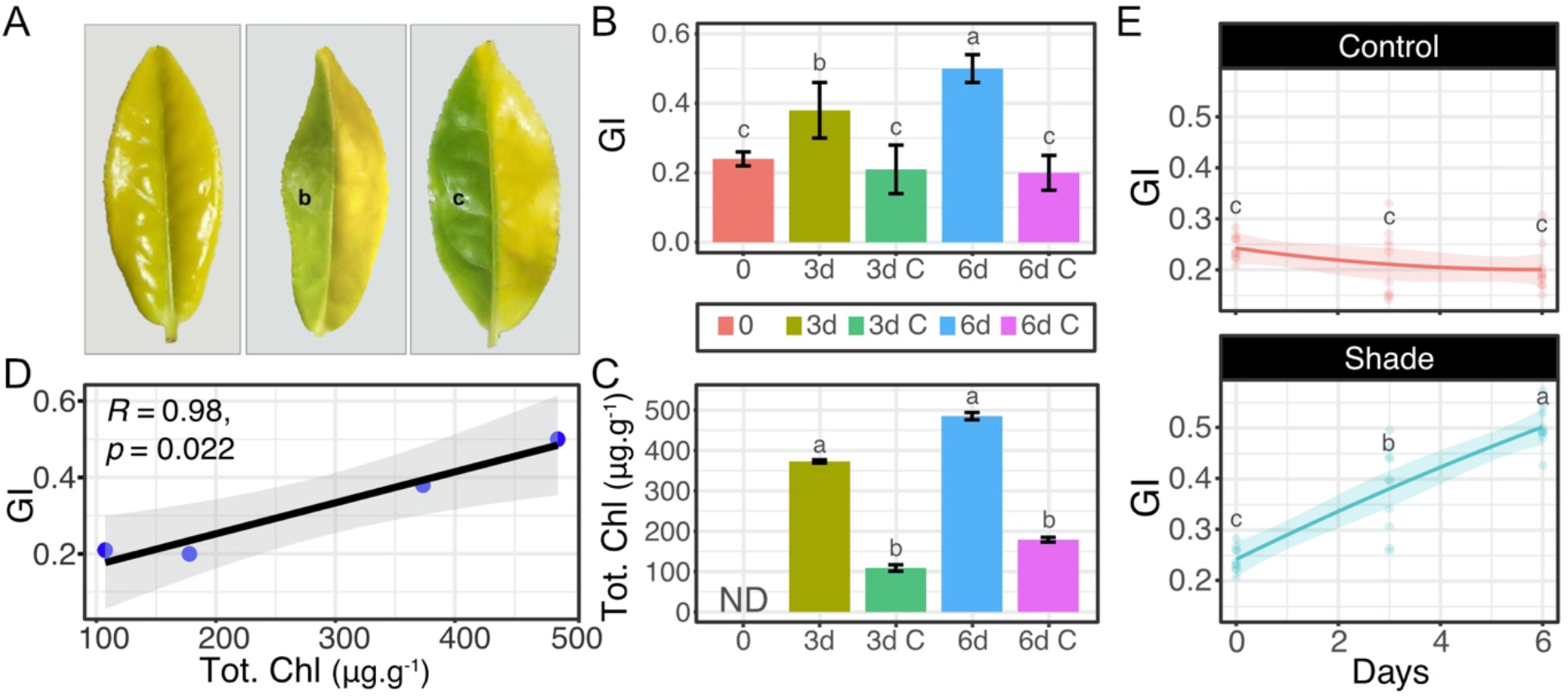
GI of leaves exposed to different light treatments. A. Leaves exposed to different light treatments: b, 3 days of shade; c, 6 days of shade; as described in (Wu et al. 2016). B. GI. C. Total chlorophyll as reported by Wu et al. (2016). D. Linear regression and Pearson correlation between GI and total chlorophyll content. E. Effect of different treatments on GI variable along the timecourse studied. Different letters indicate statistical significance differences in an ANOVA Tukey test *p* <0.05. Panel A was obtained from Wu et al. (2016).

We would suggest five different biological replicates (e.g. 5 different leaves or plants) with each of them analysed in 4 to 10 different regions for optimal results. However, we recognise this recommendation is only suitable for certain types of images. For instance, when working with developing seedlings, such as those shown in **Figure 4**, we analysed 10 different cotyledons (biological replicates) each of them determined by a single RGB value representing a 5x5 pixel area (technical replicate). This was necessary because of the size of the cotyledon in the images, the fact that they were usually homogeneous in colour, and the heterogeneity between seedlings was more relevant than technical replicates.

**Figure 4.**
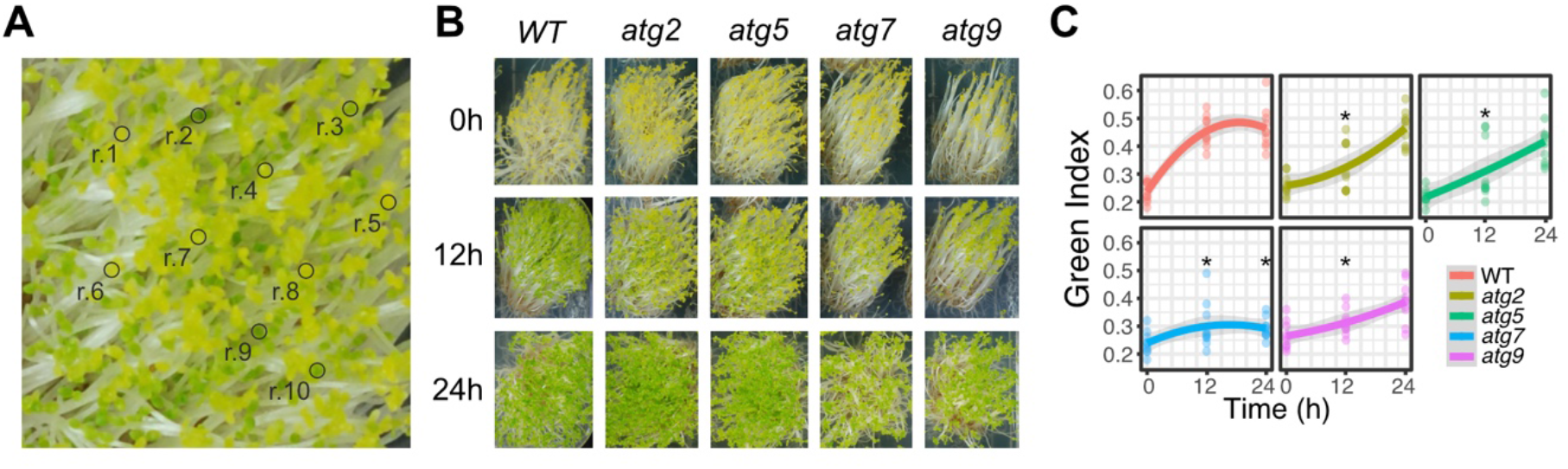
Green Index of etiolated arabidopsis seedlings exposed to light. A. Magnification of one of the genotypes tested to show the selection of replicates. B. Visual phenotype of the 5 genotypes after 0, 12 and 24h of light treatment. C. Green Index of the 5 genotypes tested at 0, 12 and 24h after light exposure (n=10). The grey regression represents +/-95% confidence level intervals for prediction for the polynomial regression. Asterisks indicate statistically significant differences (P < 0.05) by Tukey’s HSD test compared to WT at the corresponding time. Data obtained from Wijerathna-Yapa et al. (2021).

### Application of the GI method to images of biological experiments

To best illustrate the value of GI to biologists, we have considered a series of examples of biological processes where greenness changes and show the value of GI in quantifying experimental images.

### Effect of light on leaf greenness and GI

The work of Wu et al. (2016) studied the effect of light on leaves of *Camellia sinensis* L. cultivar *Baijiguan*, which had an unusual yellow leaf phenotype and when exposed to low light intensity (or shade) developed a normal green phenotype. Although there is a clear progression in green saturation and value after exposure to shade in their images, the chlorophyll assessment they reported was unable to discriminate between 3 and 6 days of shade treatment. We investigated if the GI methodology could distinguish between the clear phenotype of leaves exposed to shade and light for 6 days in images, and between the less clear phenotype of 3 and 6 days of shade treatment (**Figure 3A**). GI was able to clearly discriminate between the shade treatments and their controls at any day and between the different days of shade treatments (**Figure 3B**). Moreover, the Green Index data agreed with the total chlorophyll data (**Figure 3C**) and, despite the low number of samples, the Pearson correlation coefficient denoted a high correlation between total chlorophyll and Green Index (**Figure 3D**). Besides the bar plot, we present the data as kinetics (**Figure 3E**) to show another way of using Green Index when a time course experiment is performed.

### Effect of light on GI of etiolated seedlings

In Wijerathna-Yapa et al. (2021) we originally calculated GI to evaluate the greening process in etiolated seedlings of different genotypes. Here, the GI was calculated from different cotyledons. In **Figure 4A**, we show how heterogeneous the greening of seedlings at 12h can be, and we illustrate a putative sampling for this phenotype where a combination of green and yellow cotyledons was selected according to their abundance. As observed in **Figure 4B**, the greenness was more homogenous at 0h (mostly yellow) and at 24h (mostly green) than at 12 h, and this was clearly reflected in the Green Index (**Figure 4C**). Even though there was heterogenicity observed in greenness at 12h, the Green Index showed statistically significant differences between all the autophagy genotypes and the WT plants at 12h, but only for one genotype at 24h suggesting that most of the genotypes could reach a wildtype state by 24h (**Figure 4C**). Thus, the Green Index method was useful for establishing evidence of a statistically significant delay in greening in the autophagy genotypes tested.

### Effect of abiotic stressors on Green Index

The study by Sakuraba et al. (2014), tested the effect of mannitol, hydrogen peroxide and NaCl on WT and the autophagy mutant *atg5*. These compounds are commonly used to impose osmotic, oxidative, and saline stress respectively. The authors took pictures at 0 and 7 days after treatments. We analysed the Green Index from their images and observed that 200 mM mannitol was already enough to induce a significant difference between atg5 and WT at 7 days after treatment (**Figure 5B**), whereas no significant differences were observed with 50 mM mannitol. Likewise, 20 and 50 mM H_2_O_2_ showed significant differences between these lines in terms of Green Index, while 5 mM did not (**Figure 5B**). Finally, the lowest concentration of NaCl tested also produced significant differences to controls in terms of Green Index but not the greatest concentrations (300 and 450 mM, **Figure 5B**), which is consistent with observations of the images by eye.

**Figure 5.**
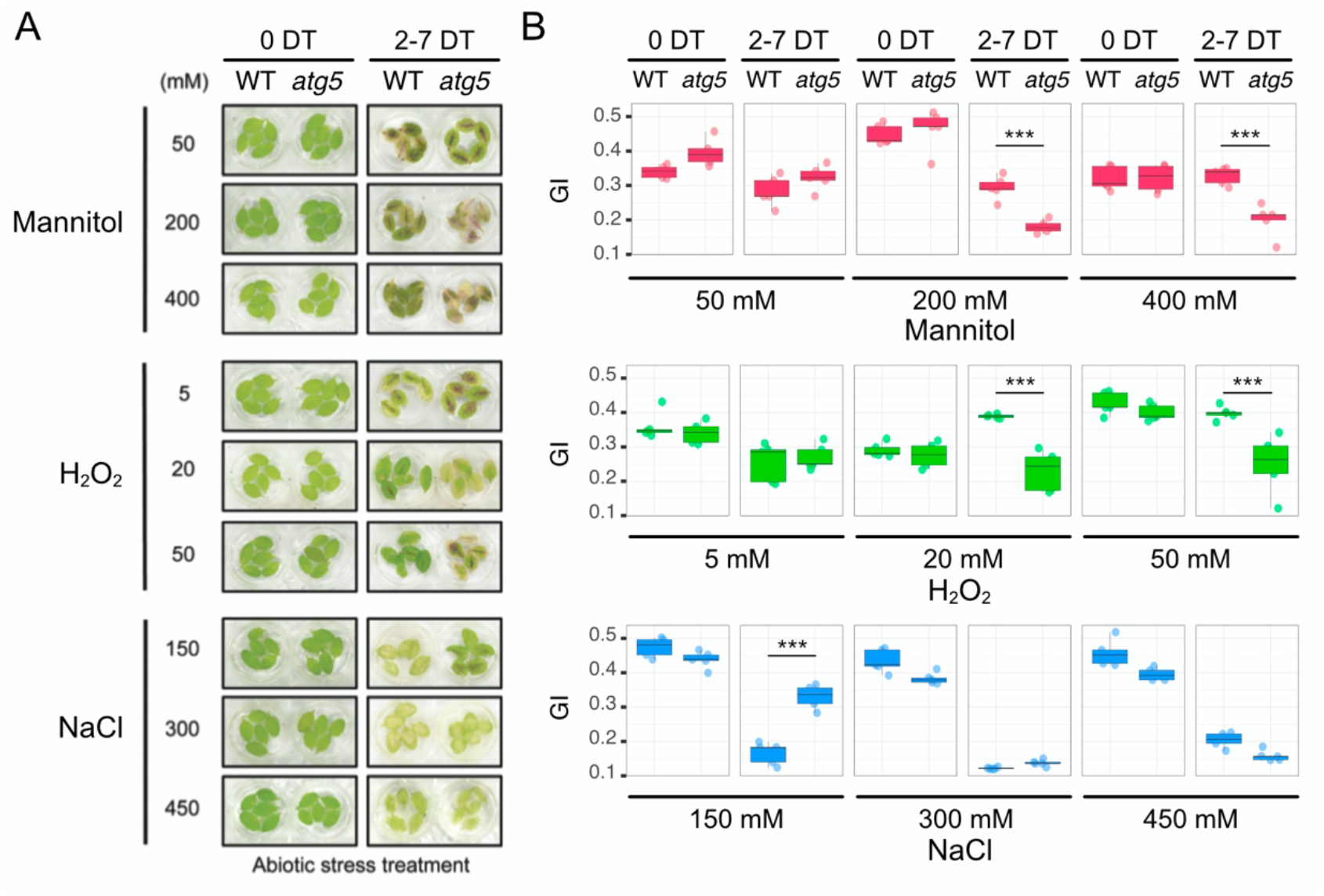
Green Index of arabidopsis leaves exposed to different types of abiotic stress. A. Phenotype of WT and atg5 leaves exposed to different concentrations of Mannitol, H_2_O_2_ and NaCl after 0 and 7 days of treatment. B. Green Index data for the samples shown in A. Asterisks indicate statistically significant differences (***, P < 0.001) by Tukey’s HSD test compared to WT. Figure in panel A was obtained from Sakuraba et al. (2014).

These results show that Green Index can be used to quantify responses under different abiotic stress conditions and its value correlated very well with the chlorophyll data provided in the same study (R=0.82, p = 3.3e-5, **Supp. Figure 2A**).

As observed in Figure 5B, the greater concentrations of NaCl resulted in severe stress for both genotypes and no significant differences were observed. The authors selected the lowest concentrations of NaCl, defined by them as mild stress, to further investigate the differences at earlier time points in the autophagy mutant (*atg5*) and the non-functional stay-green *nonyellowing1-1* (*nye1-1*). We decided to analyse these time points to see if the Green Index was able to differentiate smaller differences in phenotype. Both 3 and 5 days after treatment, Green Index values for WT were significantly different to those for *atg5*, while only five days after treatment WT was significantly different to *nye1-1* (**Figure 6**), and a strong correlation between Green Index and reported chlorophyll content was found (R=0.86, p = 0.0027, **Supp. Figure 2B**).

**Figure 6.**
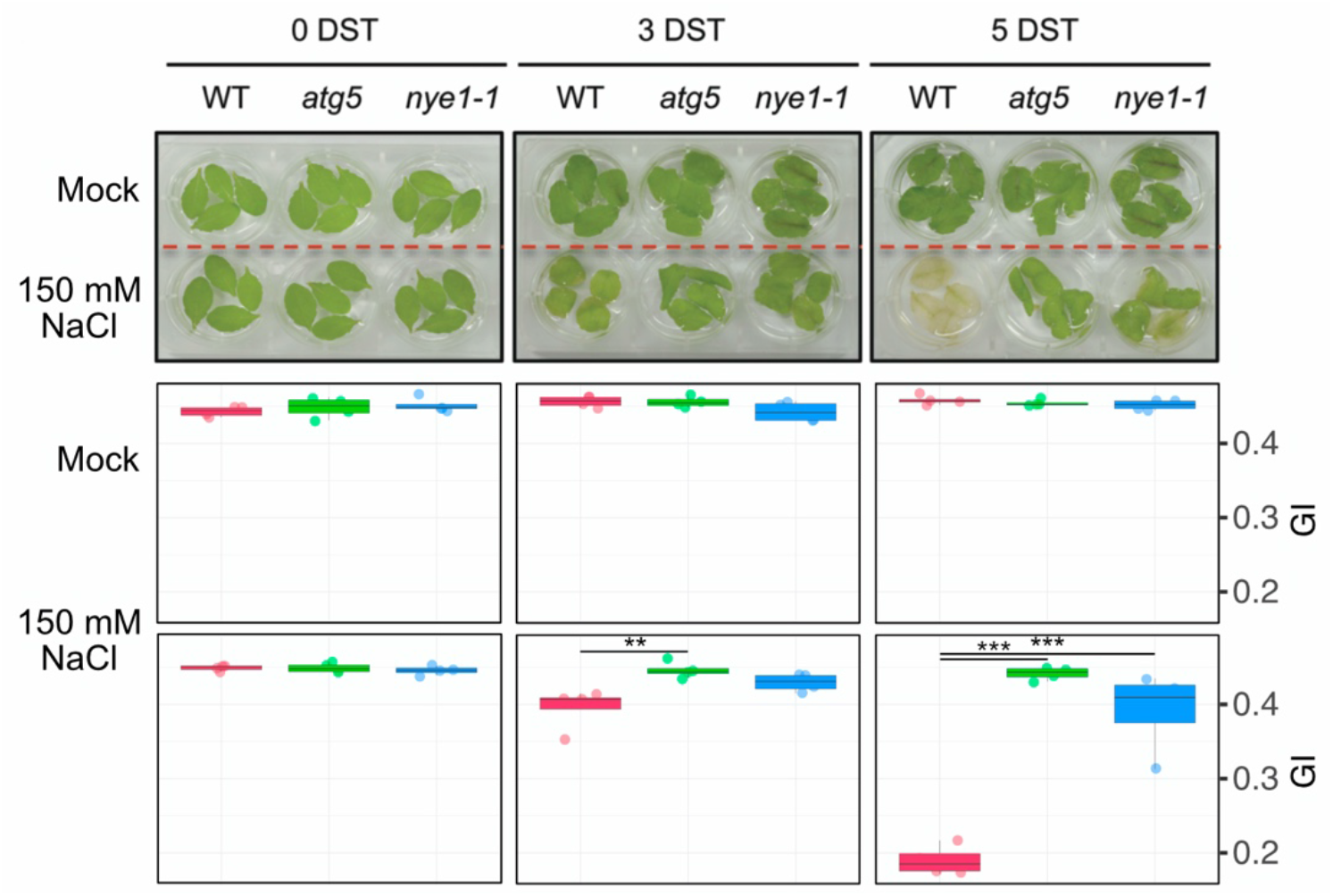
Green Index of arabidopsis leaves exposed to 150 mM NaCl. A. Phenotype of WT, *atg5* and *nye1-1* leaves exposed to NaCl 150 mM during 0, 3 and 5 days. B. Green Index data for the samples shown in A. Asterisks indicate statistically significant differences (**, P < 0.01; ***, P < 0.001) by Tukey’s HSD test compared to WT. The figure in panel A was obtained from Sakuraba et al. (2014).

This result suggests that GI could even identify subtle differences in green saturation and the value of a phenotype when different genotypes are exposed to mild abiotic stress.

### Effect of abiotic stress-induced and naturally occurring senescence on GI

We also investigated if GI could be used to assess the degree of senescence in different genotypes or leaves within the same plant. First, we investigated the GI in abiotic stress-induced senescence of WT, *atg5, nye1-1*, and the *atg5 nye1-1* double mutant (*an*) (**Figure 7**). GI showed significant differences between the WT and all lines at 5 days and all but *nye1-1* at 10 days after treatment. Again, a good correlation between GI and chlorophyll content was found (R=0.86, p = 0.00038, **Supp. Figure 2C**).

**Figure 7.**
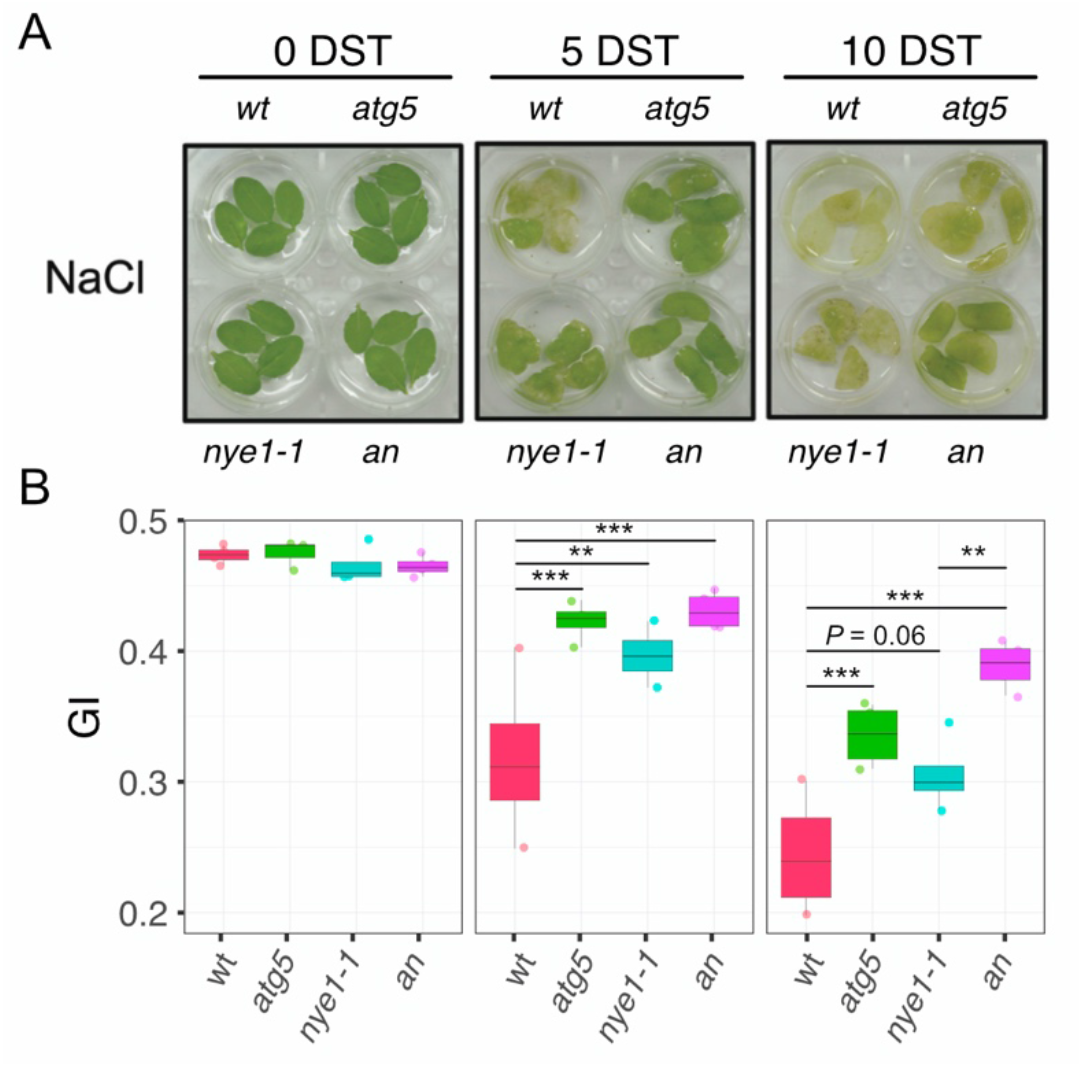
GI of arabidopsis leaves exposed to 150 mM NaCl as a means for inducing abiotic-stress dependent senescence. A. Phenotype of WT, *atg5, nye1-1* and double mutants atg5 nye1-1 (an) leaves exposed to NaCl 150 mM during 5 and 10 days. B. GI data for the samples shown in A. Asterisks indicate statistically significant differences (**, P < 0.01; ***, P < 0.001) by Tukey’s HSD test compared to WT. The figure in panel A was obtained from Sakuraba et al. (2014).

We also investigated the GI in different leaves of senescing plants at two-time points of natural senescing from the data of Koyama et al (2020) (**Figure 8A**). GIes were able to discriminate well between leaves of the same plant at the same time point (**Figure 8B**), and in many cases, the values were significantly different when comparing 65 with 75 days leaves (**Figure 8C**). Note that the study cited here was not aimed to measure greenness, thus a correct positioning of the leaves and control of light conditions could enhance the differences observed by GI.

**Figure 8.**
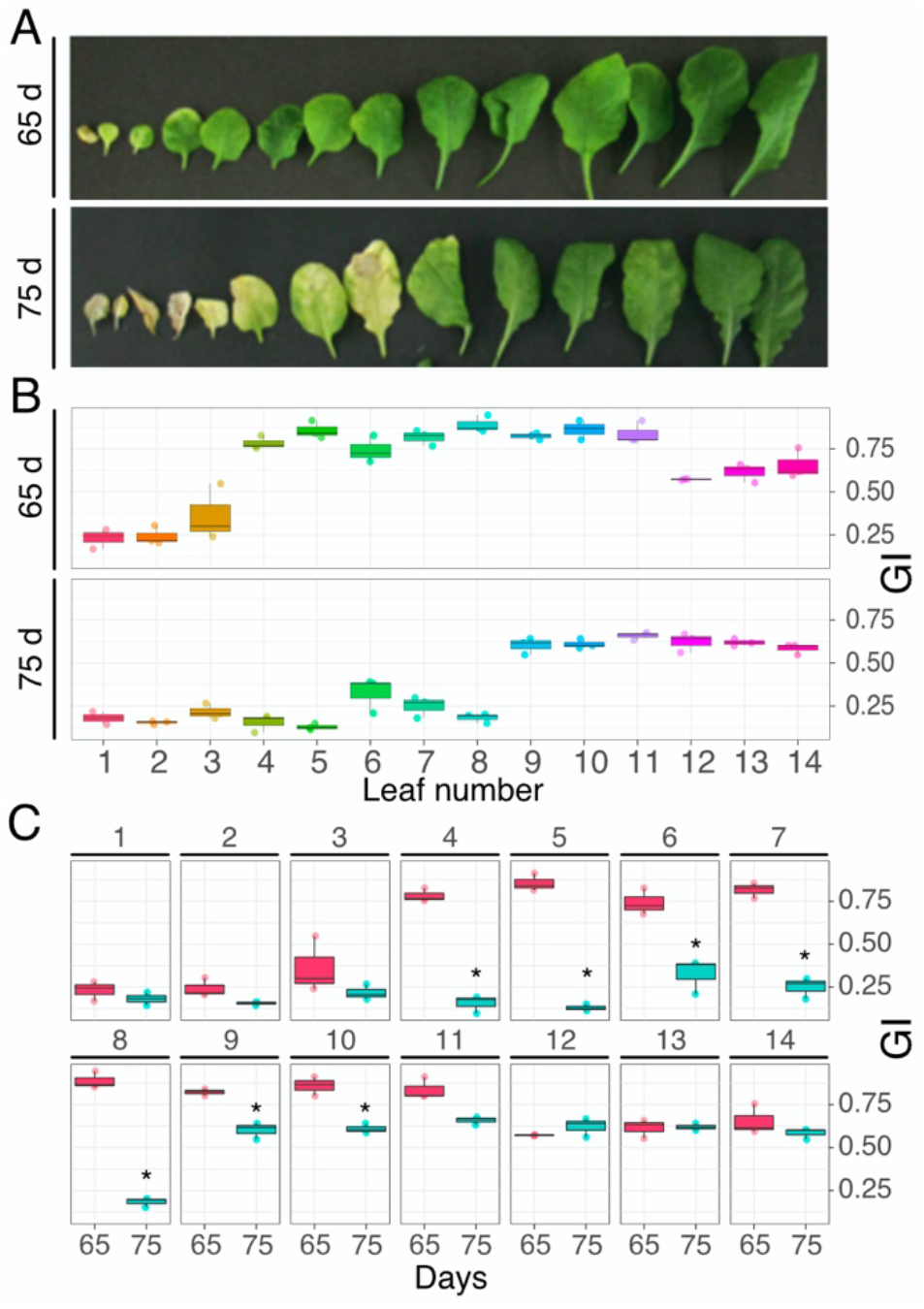
GI of arabidopsis leaves during senescence. A. Phenotype of 65-da and 75-day old arabidopsis leaves. B. GI data grouped by age. C. GI data grouped by leaf number. Asterisks indicate statistically significant differences by Tukey’s HSD test. The figure in panel A was obtained from Koyama et al (2020).

To further explore the potential of GI in studies of senescence, we compared the GI scores of leaves of the same age for different genotypes and treatments. In the study from Kim et al. (2015), the phenotype of senescing leaves of arabidopsis Columbia (Col) and Wassilewskija (WS) ecotypes, together with *dab4-1* (a jasmonic acid receptor mutant), and *ein2-1* (an ethylene insensitive mutant) were analysed upon exposure to air, methyl jasmonate (meJA) and ethylene. Under air conditions, the GI were not statistically different, however, under ethylene and meJA, significant differences were observed among the genotypes (**Figure 9**).

**Figure 9.**
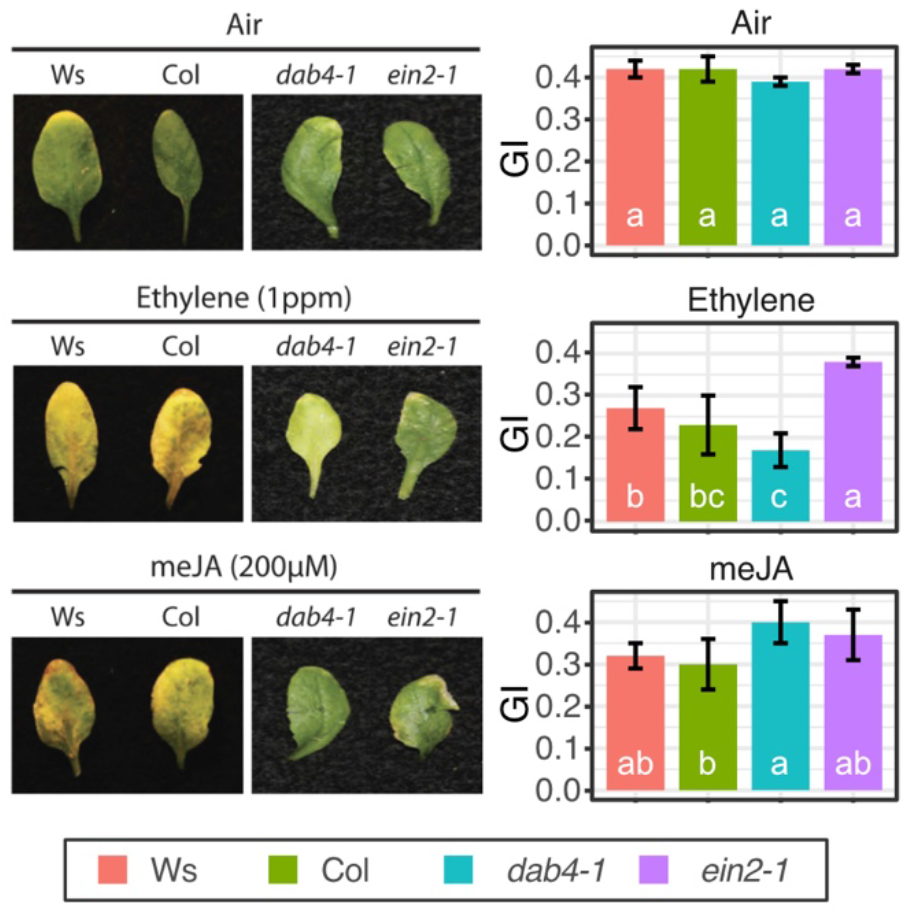
GI of arabidopsis leaves of different genotypes during senescence upon exposure to air, ethylene and meJA. Phenotype of Columbia (Col), Wassilewskija (WS), *dab4-1* (Jasmonic Acid receptor mutant), and *ein2-1* (ethylene insensitive mutant) exposed to air, ethylene and meJA, obtained from Kim et al (2015), and their respective GI values. Different letters indicate statistically significant differences by Tukey’s HSD test.

### GI to evaluate nutritional stress in seedlings

Excess or deficiency of nutrients can be a cause of plant stress, thus impacting plant phenotype. Liang et al. (2017) phenotyped arabidopsis plants in response to different concentrations of glucose and nitrogen. Using those images, we determined GI to observe if it was sensitive enough to quantify the mild differences in green saturation and value of leaves from different nutrients treatments. Indeed, the nutrient replete leaves score GI values up to 0.43, whereas the most nutrient depleted plants showed values as low as 0.26 (**Figure 10A**). The GI results also agreed with the estimated chlorophyll content determined in the original publication for the different concentrations of both glucose and nitrogen (**Figure 10B-C**). For most concentrations of nitrogen, increasing glucose concentration showed a positively effect on greenness and GI value; however, at 0.1 mM KNO_3_, the greatest concentrations of glucose were detrimental to greenness (**Figure 10D**). Nitrogen increased the greenness and GI values, and such an increase grew with increasing concentrations of glucose (**Figure 10D**). Finally, we correlated the GI values with the estimated chlorophyll content in the original publication observing that there was a significant positive correlation between these parameters (**Figure 10G**).

**Figure 10.**
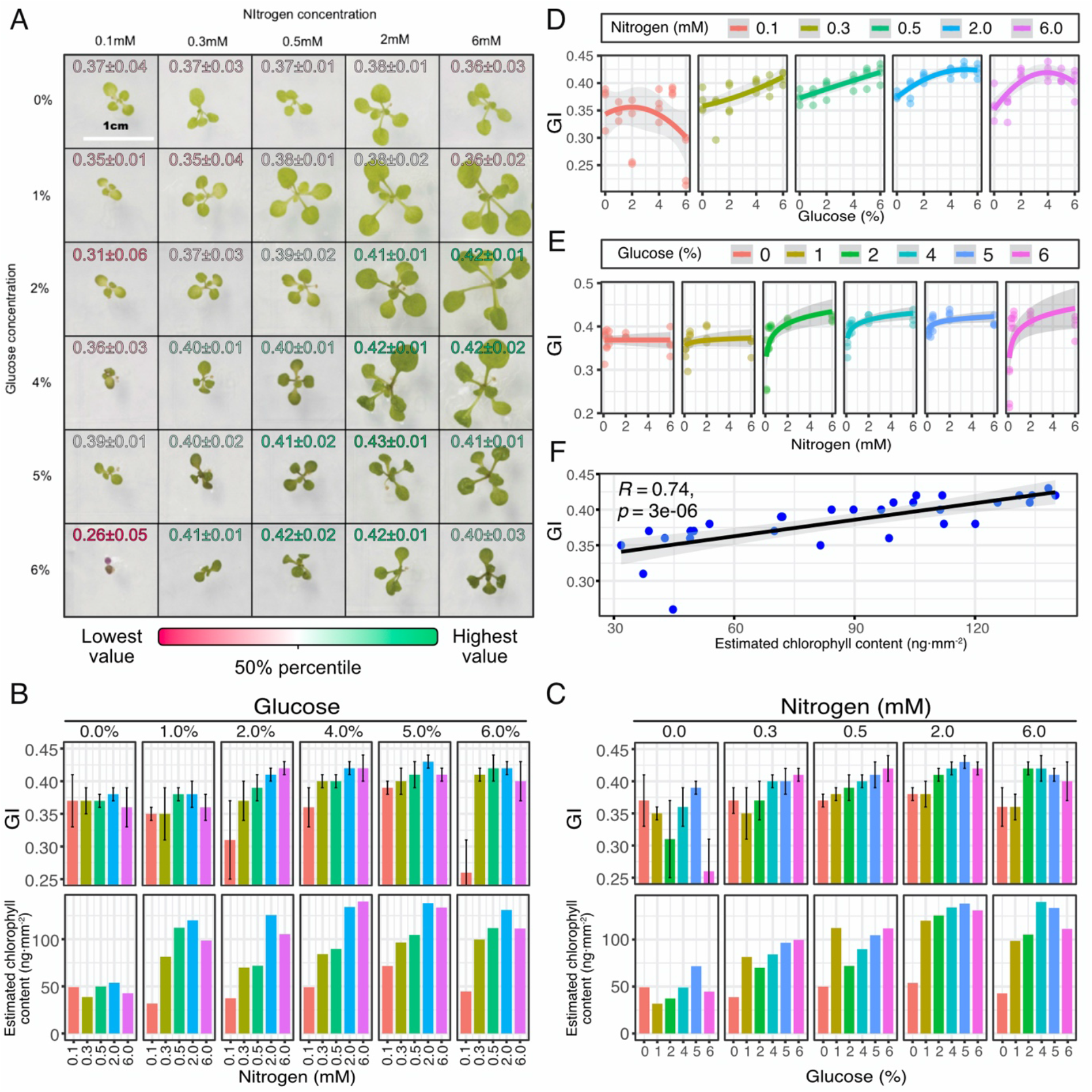
GI analysis of arabidopsis seedlings exposed to different concentrations of glucose and KNO_3_. **A**. Plant phenotypes and GI values (image taken from Liang et al. (2017)). **B** and **C**. Bar chart of GI values and the estimated chlorophyll content determined by Liang et al. (2017) for different glucose and nitrogen concentrations. T-bars indicate the standard deviation. **D** and **E**. Scatter plots and trend for GI values relative to different glucose and nitrogen concentrations. **F**. Correlation between GI values and the estimated chlorophyll content determined by Liang et al. (2017).

### GI to evaluate chemical treatment for post-harvest preservation

Meitha et al. (2022) tested the effect of chitosan on the post-harvest preservation of spinach leaves and determined chlorophyll content as one of the parameters to indicate leaf health (**Figure 11**). GI data was able to discriminate between the days of post-harvest and between the treatments (**Figure 11B**). The GI values showed a very good correlation with chlorophyll content (R¿0.94, p=0.00049, **Figure 11D**).

**Figure 11.**
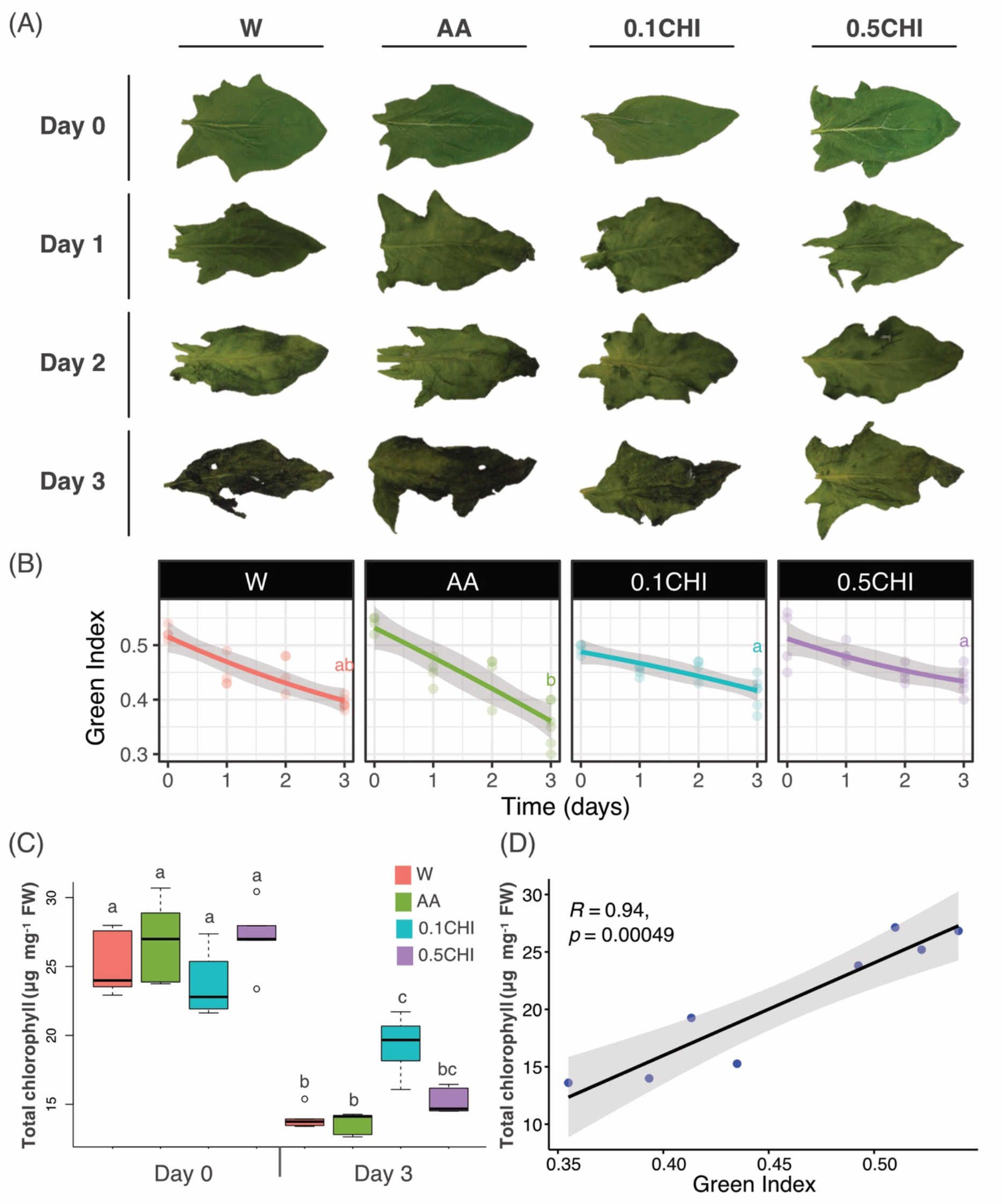
GI analysis of spinach leaves treated with acetic acid and chitosan. Taken from Meitha et al. (2022). **A**. Phenotype of leaves. **B**. GI. **C**. Chlorophyll content. **D**. Correlation between chlorophylls and GI. AA, 1% acetic acid (v/v); 0.1CHI, 0.1% chitosan (w/v); 0.5CHI, 0.5% chitosan (w/v); W, water.

### Comparison of GI values obtained with an SLR vs smartphone

Given that different cameras have different sensors that are sensitive to different wavelengths of light, they can give different RGB values for the same object and light condition. Moreover, the algorithms for processing the raw data captured by the sensor that is used to produce the image (i.e. the RGB values), as well as the settings (e.g. exposure time, aperture size, ISO sensitivity, etc.) also vary between cameras, resulting in different RGB values. To test the effect of these differences on GI we compared values from a professional single-lens reflex mirrorless camera (Nikon D7000 digital SLR camera) and a mirror-based phone camera (iPhone Xs). The images obtained by these devices looked different to the eye, being more saturated in colour from the phone camera than the SLR camera (**Figure 12**). The GI was also different from to data obtained with each camera, however, the trend and difference between time points was constant (**Figure 12**). So, while GI of images obtained from these different styles of cameras are not a constant and cannot be directly compared, any camera is likely to be useful to determine GI for a particular experiment.

**Figure 12.**
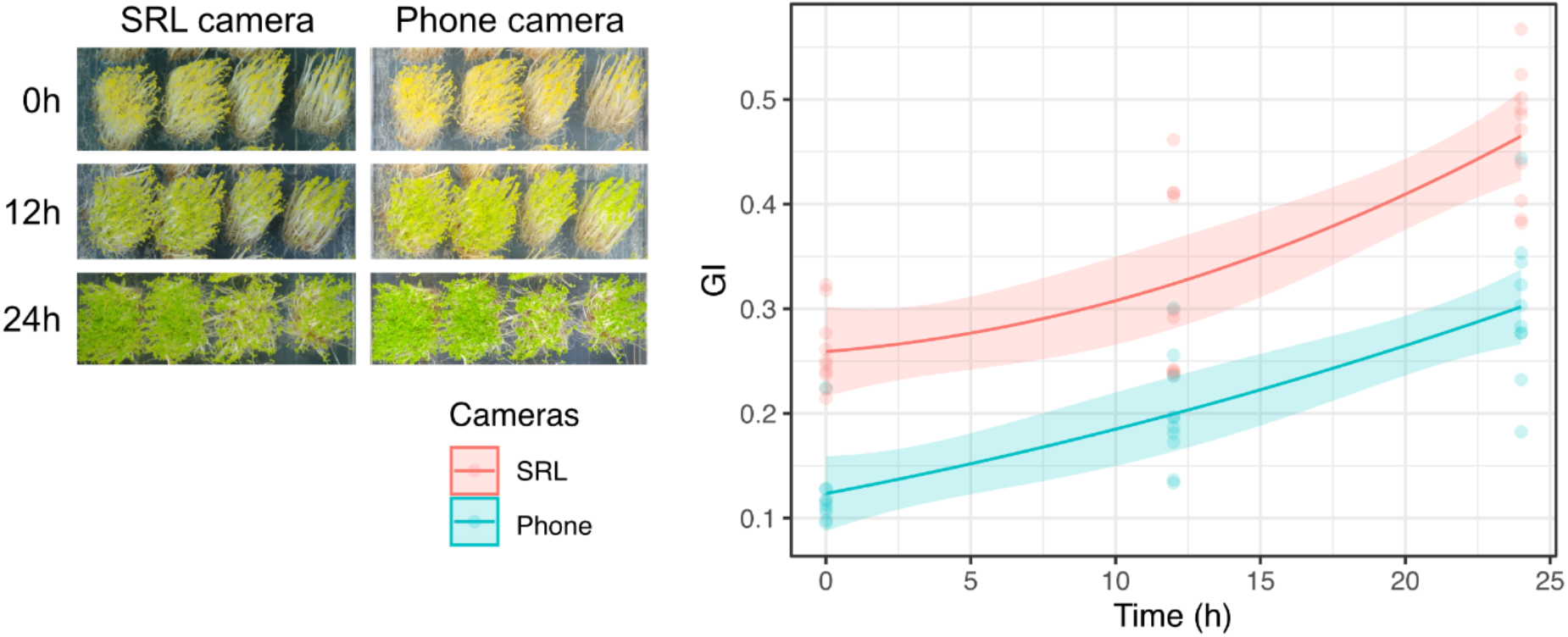
Comparison of GI calculated from pictures taken by a single-lens reflex (SLR) mirrorless camera and a mirror-based phone camera. The image shows de-etiolated arabidopsis *sugar dependent mutant* (*sdp1*) seedlings exposed to light for 0, 12 and 24h. The graph represents the GI at 0, 12 and 24 h calculated from the SRL and phone camera images.

### Summary of procedures to use and calculate GI

In the above sections, we have shown that GI calculations can be useful to compare plants based on greenness, either to compare different genotypes in a given treatment, to compare the response of a genotype to different treatments (e.g. abiotic, nutritional stress), or different developmental stages. We also showed that GI correlates well with chlorophyll content, meaning that GI could be used with caution to infer chlorophyll content in different samples based on a simple picture taken with a phone camera. Given that chlorophyll extraction and quantification require reagents (e.g. organic solvents) equipment (e.g. spectrophotometer) and a laboratory, as well as time, this technique has limited accessibility. However, with over 16 billion mobile phones operating in the world in 2023, being 6.9 billion smartphones (Statista 2023), the ability to capture GI is widely available.

In **Figure 13**, we illustrate a summary of the procedure to use GI.

**Figure 13.**
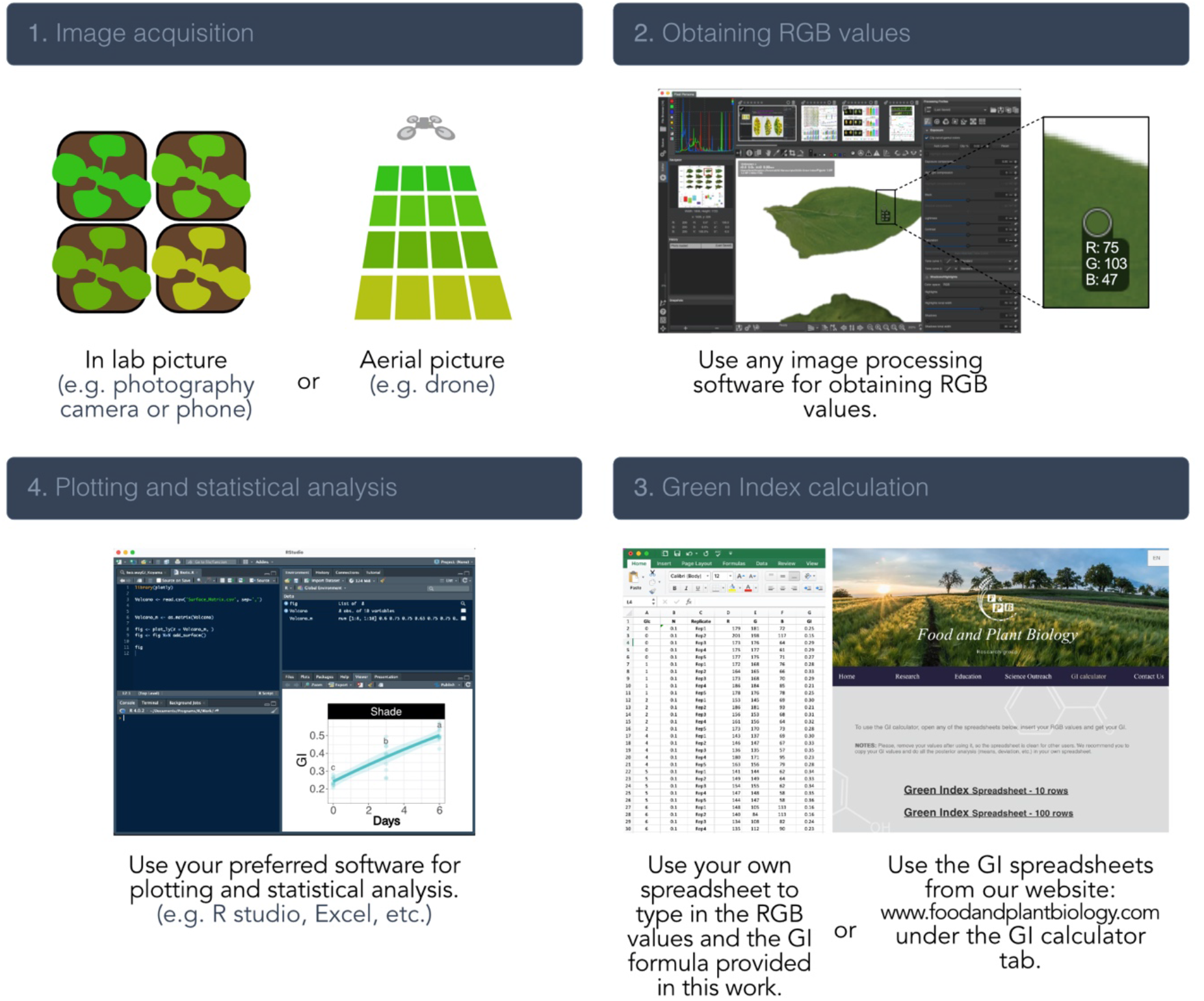
Highly accessible summarized procedure to determine GI. 1. Image acquisition. The input required for GI determination is any image from which RGB values can be extracted. 2. Obtaining RGB values. Each pixel in a picture has RGB values that can be obtained by multiple image processing software. In this study, we used the open software RawTherapee. For this program, the mean RGB values for the defined area are presented (not the RGB of a single pixel). 3. GI calculation. Either use the formula provided in this study to determine GI on a spreadsheet (eq. 6) or use the GI calculator found at www.foodandplantbiology.com. 4. Plotting and statistical analysis. Using the GI obtained for your study, replicates and treatment, proceed to represent them as preferred and look for statistically significant differences among your samples/treatments.

To use GI as a proxy for chlorophyll estimation, it is important to bear in mind that GIes are affected by lighting conditions. Therefore, we recommend that in outdoor settings the lighting is similar in the whole frame and the data be presented as relative values.

## Conclusions

We have developed GI as a parameter useful to discriminate different levels of greenness in images of leaves. The calculation of this parameter is based on RGB values that can be easily obtained from pictures, it does not require any informatic skills, and it is accessible to anyone with a computer, free software, and a camera. We have shown that the differences in GI are useful to quantitatively discriminate between healthy and damaged leaves, leaves at different stages of development, and in analysis of abiotic stress. Moreover, we show GI correlates well with leaf chlorophyll content, providing an easy and low-cost alternative to chlorophyll determination. This approach to assessing greenness is widely accessible to scientists regardless of funding or equipment, when non-destructive chlorophyll content is needed, or when limited amount of tissue is available (e.g. 5-days-old arabidopsis seedlings) or rapid large-scale assessment is required (e.g. canopy of a large field). In the future, the incorporation of this formula in image-analysis software and its automatization would further improve its usage.

## Acknowledgment

Figure 2, 3 reproduce parts of figure 1 from Wu et al. (2016) which is free to be reproduced under the Creative Commons Attribution 4.0 International License (http://creativecommons.org/licenses/by/4.0/). Figure 4 reproduces parts of figure 2 from Wijerathna-Yapa et al. (2021) which is free to be reproduced under the Creative Commons Attribution 4.0 International License (http://creativecommons.org/licenses/by/4.0/). Figure 5, 6 and 7 reproduce parts of figure 1, 2 and 5 in Sakuraba et al. (2014) which are free to be reproduced under the Creative Commons Attribution License (http://creativecommons.org/licenses/by/3.0/). Figure 8 reproduces parts of figure 7 from Koyama et al. (2017) which is free to be reproduced under the Creative Commons Attribution 4.0 International License (http://creativecommons.org/licenses/by/4.0/). Figure 9 reproduces parts of figure 1 from Kim et al. (2015) which is free to be reproduced under the Creative Commons Attribution 4.0 International License (http://creativecommons.org/licenses/by/4.0/). Figure 10A in our manuscript reproduces figure 3 in Liang et al. (2017) which is free to be reproduced under the Creative Commons Attribution 4.0 International License (http://creativecommons.org/licenses/by/4.0/). Figure 11A-C in our manuscript reproduces figure 1A,B,D in Meitha et al. (2022) for which we got permission to be republished in BioRxiv under Licence number 5613801423598 and license date Aug 21, 2023, granted by Springer Nature.

## Notes

### Competing Interest Statement

The authors have declared no competing interest.

