## Supplemental data for "Green Index: a widely accessible method to quantify greenness of photosynthetic organisms"

A

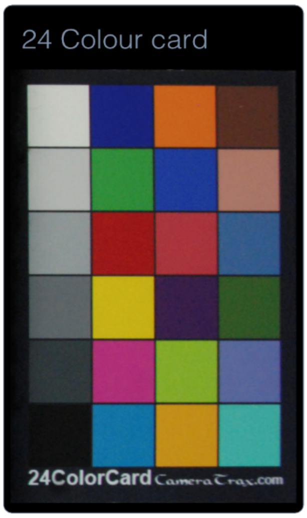


B

| Colour | R | G | B | pre-GI_a | pre-GI_b | pre-GI_c | GI |
|---|---|---|---|---|---|---|---|
| Turquoise | 66 | 169 | 159 | 0.616 | 0.799 | 3.97 | 0.331 |
| Orange-yellow | 196 | 144 | 28 | 0.650 | 0.753 | 3.05 | 0.254 |
| French blue | 4 | 164 | 116 | 0.759 | 0.852 | 5.77 | 0.481 |
| Black | 17 | 19 | 18 | 0.848 | 0.757 | 3.11 | 0.259 |
| Sky-blue | 91 | 103 | 155 | 0.675 | 0.695 | 2.28 | 0.190 |
| Lime | 133 | 170 | 45 | 0.676 | 0.859 | 6.08 | 0.507 |
| Violet | 176 | 52 | 119 | 0.552 | 0.565 | 1.30 | 0.108 |
| Darkest grey | 48 | 56 | 58 | 0.804 | 0.817 | 4.46 | 0.372 |
| **Dark green** | **45** | **86** | **34** | **0.878** | **0.882** | **7.50** | **0.625** |
| Purple | 56 | 24 | 71 | 0.735 | 0.748 | 2.96 | 0.247 |
| Yellow | 200 | 178 | 16 | 0.616 | 0.742 | 2.88 | 0.240 |
| Dark grey | 95 | 101 | 101 | 0.742 | 0.758 | 3.14 | 0.261 |
| Blue | 47 | 91 | 138 | 0.746 | 0.759 | 3.16 | 0.263 |
| Pink | 171 | 52 | 61 | 0.634 | 0.647 | 1.83 | 0.153 |
| Red | 158 | 27 | 21 | 0.671 | 0.641 | 1.78 | 0.148 |
| Grey | 142 | 151 | 150 | 0.552 | 0.698 | 2.31 | 0.193 |
| Coral | 171 | 120 | 103 | 0.616 | 0.681 | 2.14 | 0.178 |
| Blue 2 | 22 | 61 | 154 | 0.719 | 0.692 | 2.24 | 0.187 |
| **Green** | **51** | **141** | **60** | **0.801** | **0.922** | **11.8** | **0.979** |
| Light grey | 178 | 180 | 177 | 0.431 | 0.614 | 1.59 | 0.133 |
| Brown | 86 | 48 | 35 | 0.774 | 0.780 | 3.55 | 0.296 |
| Orange | 200 | 99 | 21 | 0.710 | 0.680 | 2.12 | 0.177 |
| Navy | 4 | 31 | 120 | 0.748 | 0.673 | 2.06 | 0.172 |
| Lightest grey | 196 | 199 | 192 | 0.363 | 0.546 | 1.20 | 0.100 |
| MIN | | | | 0.363 | 0.546 | 1.20 | 0.100 |
| | | | | 0.621 | 0.734 | 6.48 | 0.540 |
| MAX | | | | 0.878 | 0.922 | 11.8 | 0.979 |

C

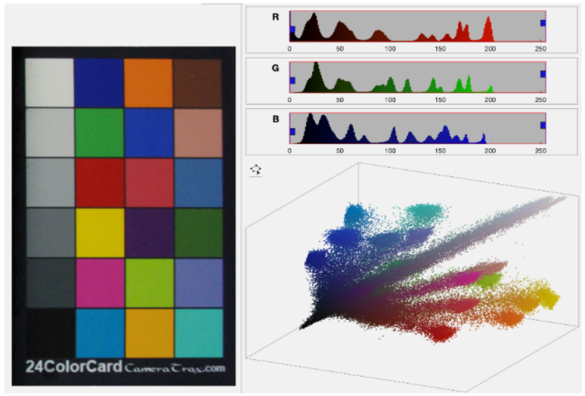


D

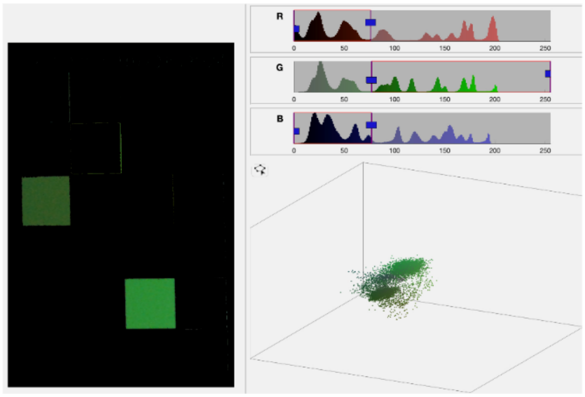


**Supplemental figure 1.** Analysis of greenness in a 24-colour card for de optimization of our GI formula. **A**. 24 colour card used in this study. **B**. RGB values for each colour and results obtained with the different formulas. **C**. Distribution of pixels for the 24-colour card without filtering. **D**. Pixels remining after doing the colour segmentation based on pixels that are between 0-75 for R, 75-255 for G, and 0 to 75 for B.

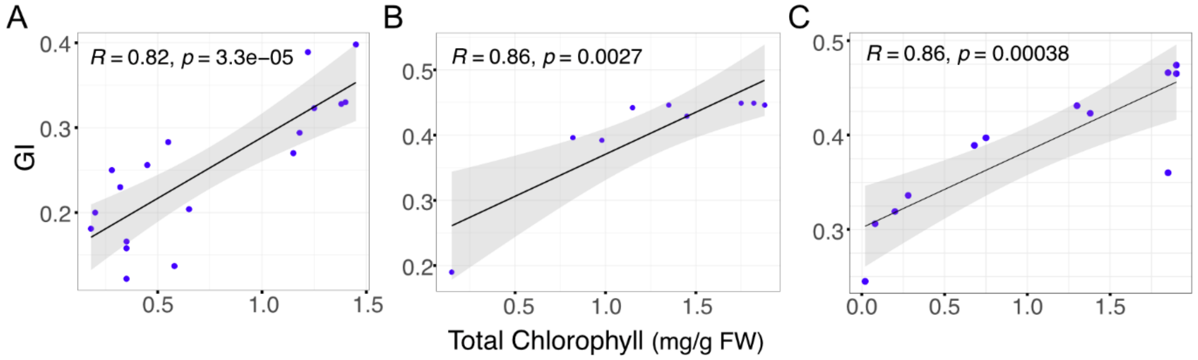


**Supplemental figure 2.** Pearson correlation between GI and total chlorophyll content. **A**. Correlation for data obtained by incubating leaves with different abiotic stressors (mannitol, hydrogen peroxide, and sodium chloride, Fig. 1 in Sakuraba et al. 2014). **B**. Correlation for data obtained by incubating leaves with 150 mM sodium chloride for 0, 3 and 5 days, Fig. 2 in Sakuraba et al. 2014. **C**. Correlation for data obtained by incubating leaves with 150 mM sodium chloride for 0, 5 and 10 days, Fig. 5 in Sakuraba et al. 2014. Blue dots represent independent samples with a regression curve applied (in black) and 95 % confidence intervals (in grey shading).
